# Profiling microRNA expression during senescence and aging: mining for a diagnostic tool of senescent-cell burden

**DOI:** 10.1101/2024.04.10.588794

**Authors:** Moritz Weigl, Teresa L. Krammer, Marianne Pultar, Matthias Wieser, Selim Chaib, Masayoshi Suda, Andreas Diendorfer, Kseniya Khamina-Kotisch, Nino Giorgadze, Tamar Pirtskhalava, Kurt O Johnson, Christina L. Inman, Ailing Xue, Ingo Lämmermann, Barbara Meixner, Lichao Wang, Ming Xu, Regina Grillari, Mikolaj Ogrodnik, Tamar Tchkonia, Matthias Hackl, James L Kirkland, Johannes Grillari

**Affiliations:** Ludwig Boltzmann Institute for Traumatology, The Research Center in Cooperation with AUVA, 1200 Vienna, Austria; Austrian Cluster for Tissue Regeneration, 1200 Vienna, Austria; Department of Physiology and Biomedical Engineering, Mayo Clinic, Rochester, MN, USA; Division of General Internal Medicine, Mayo Clinic, Rochester, MN, USA; TAmiRNA GmbH, Vienna, Austria; Evercyte GmbH, Vienna, Austria; Ludwig Boltzmann Research Group Senescence and Healing of Wounds, Vienna, Austria; Rockfish Bio AG, Vienna, Austria; UConn Center on Aging, UConn Health, Farmington, CT, USA; Department of Genetics and Genome Sciences, UConn Health, Farmington, CT, USA; Department of Cardiovascular Biology and Medicine, Juntendo University Graduate School of Medicine, 3-1-3 Hongo, Bunkyo-ku, Tokyo 113-8421, Japan; Institute for Molecular Biotechnology, BOKU University, Vienna, Austria

## Abstract

In the last decade cellular senescence, a hallmark of aging, has come into focus for pharmacologically targeting aging processes. Senolytics are one of these interventive strategies that have advanced into clinical trials, creating an unmet need for minimally invasive biomarkers of senescent cell load to identify patients at need for senotherapy. We created a landscape of miRNA and mRNA expression in five human cell types induced to senescence *in-vitro* and provide proof-of-principle evidence that miRNA expression can track senescence burden dynamically *in-vivo* using transgenic p21^high^ senescent cell clearance in HFD fed mice. Finally, we profiled miRNA expression in seven different tissues, total plasma, and plasma derived EVs of young and 25 months old mice. In a systematic analysis, we identified 22 candidate senomiRs with potential to serve as circulating biomarkers of senescence not only in rodents, but also in upcoming human clinical senolytic trials.

## Main

Significant progress in understanding diseases related to aging has been made over the last few years by identifying senescent cells as common substrate of a plethora of age-related diseases^1,2^ termed senopathies^3^. Since the initial discovery of senolytics^4^, research on senescent cell targeting compounds progressed quickly^5–14^ and clinical trials of this new class of drugs are ongoing^15^. This creates a necessity for companion diagnostics of senescence that allow early assessment of safety and efficacy of senolytic drug candidates and the detection of sensitive clinical endpoints^16^. DNA methylation based epigenetic aging clocks were recently shown to lack sensitivity to the senescent phenotype^17^, and previous efforts have largely focused on the protein-component of the senescence-associated secretory phenotype (SASP)^18,19^ that has an unknown potential overlap to non-senescent inflammatory conditions and thus might be sensitive but not specific. In contrast, miRNAs of the miR-17-92 cluster turned out as commonly downregulated in various models of senescence and aging^20^. In this study we therefore aim to address the prevailing lack of a systematic exploration of the potential of non-coding microRNAs to serve as a biomarker of senescence.

MicroRNAs are important post-transcriptional regulators of gene expression. Their presence in biofluids such as plasma or extracellular vesicles (EVs) can reflect transcriptional changes in tissues such as bone^21,22^, indicating that the non-coding transcriptome could serve as an abundant source of novel biomarkers^23^. Moreover, we and others have previously proposed extracellular miRNAs as being part of the SASP^24–27^, and that miRNAs released from senescent cells could play an underappreciated role in causing detrimental effects to tissue homeostasis during aging^28^.

In this study, we analyzed miRNA and mRNA expression in five different human cell types induced to senescence and identified common and unique gene expression features regulated during senescence. We provide proof-of-principle evidence that miRNA expression patterns can follow differential senescent-cell abundance in bulk tissues by taking advantage of *PLD* mice, a novel transgenic model that allows selective ablation of p21^high^ senescent cells. Finally, we systematically identified a set of potential *senomiRs*, microRNA biomarkers of senescence, by profiling miRNA expression in aged (n=7) *vs*. young (n=5) mice in seven different tissues and two fractions of the circulation, total plasma and plasma derived EVs, respectively. By correlating tissue expression of miRNAs with the established senescence markers p16 and p21 and subsequently correlating levels of miRNAs in tissues with levels detected in the circulation, we propose a selection of 22 circulating *senomiRs* as a potential minimally invasive biomarker of organismal senescent cell load.

## Results

### mRNA and miRNA expression analysis during cellular senescence reveals few common but predominantly cell-type specific patterns

To investigate changes in miRNA expression during cellular senescence, RNA from cells was collected three weeks after induction of stress-induced senescence (SIPS) using Doxorubicin (DOXO) *vs*. quiescent (QUI) cells as controls. As the senescent phenotype is cell-type dependent, five different human primary cell types - skin fibroblasts (HDFs), microvascular endothelial cells from skin (HDMVECs), umbilical vein endothelial cells (HUVECs), renal proximal tubular epithelial cells (RPTECs), and adipose derived mesenchymal stem cells (ASCs) from three different donors each were used (Figure 1a). To connect differential miRNA expression to mRNA expression changes, we performed both RNA and small RNA sequencing.

**Figure 1:**
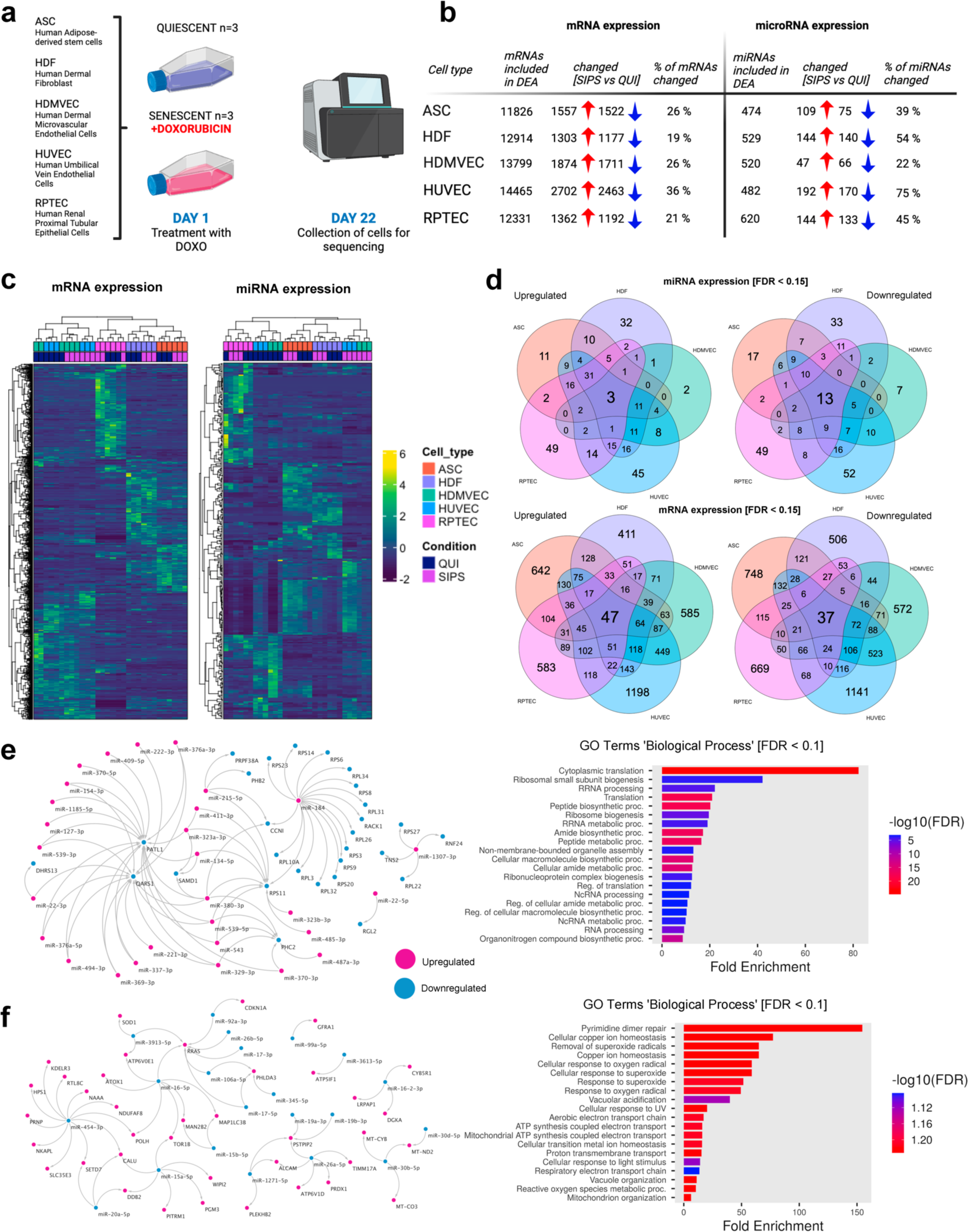
Senescent cells from different tissues have distinct mRNA and microRNA expression profiles with a core set of mRNAs and microRNAs shared among different senescent cell types. **a,** Experimental scheme for the *in-vitro* induction of senescence using Doxorubicin (DOXO) in five different cell types. **b,** Differential expression results of mRNA and microRNA profiling in five different human primary cell types. The table lists the number of mRNAs and microRNAs, respectively, which were included for DEA and that were differentially up- or downregulated (FDR < 0.15). **c,** Unsupervised heatmap clustering using RPM values for mRNA and microRNA expression. The top 3000 mRNAs and top 300 microRNAs according to CV% were used to prepare heatmaps. Pearson correlation was used for clustering of samples (columns) and mRNAs/microRNAs (rows). **d,** Differentially expressed mRNAs and microRNAs were compared between cell types and used for VENN analysis. **e – f,** mRNAs and microRNAs commonly regulated in four out of five senescent cell types were used for creating correlation networks, microRNA-mRNA pairs correlated with an adjusted p value < 0.01 were used for network construction. **e,** The upper network shows correlations between upregulated microRNAs and downregulated mRNAs, the lower network in **f,** shows correlations between downregulated microRNAs and upregulated mRNAs. For both analyses, all mRNAs in the respective networks were used for GO term enrichment using the category ‘Biological Process’. The Top 20 terms (FDR < 0.1) are shown.

The senescent phenotype was confirmed by cell morphology and senescence-associated β-galactosidase staining (Extended Data Fig. 1a), along with the detection of a significant (FDR < 0.1) increase in *CDKN1A* (p21) expression using RNA-sequencing in all senescent cell types, CDKN2A (p16) in HDMVECS and HUVECs, and lamin B1 decrease in all but ASCs and HDFs (Extended Data Fig. 1b). Cell-cycle arrest was confirmed by lack of BrdU incorporation in four cell types (Extended Data Fig. 1c). The log_2_ fold changes of different senescence markers were compared in Extended Data Fig. 1d.

Between 11826 (ASC) and 14465 (HUVEC) distinct mRNAs in each cell type were included for differential expression analysis (DEA) with 19 % (HDF) to 36 % (HUVEC) of mRNAs showing a significant up- or downregulation following the induction of senescence as assessed by an adjusted p-value (FDR) below 0.15. Small RNA-seq included between 474 (ASC) and 620 (RPTEC) miRNAs for DEA in each cell type with 22 % (RPTEC) to 75 % (HUVEC) of miRNAs showing a significant up- or downregulation during senescence (FDR < 0.15) (Figure 1b & Extended Data Figure 3).

To compare mRNA and miRNA expression, unsupervised heatmap clustering analysis was performed (Figure 1c). RPM values of the top variable 3000 mRNAs and top 300 miRNAs according to coefficient of variation (CV%) were included in heatmaps, revealing a high degree of senescent cell-type specificity of mRNA and miRNA expression profiles (Figure 1c).

Comparison of mRNA expression changes revealed common and unique features regulated in the different senescent cell types. 47 and 37 mRNAs (Supplementary Tables 1 & 2) were found up- or downregulated (FDR < 0.15) in all five senescent cell types (Figure 1d). However, most differentially expressed mRNAs were unique, with 642 (ASC), 411 (HDF), 585 (HDMVEC), 1198 (HUVEC), and 583 (RPTEC) mRNAs only upregulated (FDR < 0.15) in a single senescent cell type. Similarly, the largest fraction of mRNAs was uniquely downregulated (FDR < 0.15), with 748 (ASC), 506 (HDF), 572 (HDMVEC), 1141 (HUVEC) and 669 (RPTEC) mRNAs downregulated only in a single senescent cell type (Figure 1d).

Changes of miRNA expression detected in different senescent cell types showed parallel results with 3 (miR-215-5p, miR-203a-3p, miR-222-3p) up-, and 13 (miR-99a-3p/5p, miR-26a/b-5p, miR-101-3p, miR-30b/d-5p, miR-424-3p, miR-19a/b-3p, miR-16-5p, miR-16-2-3p, miR-92a-3p) miRNAs downregulated (FDR < 0.15) in all five senescent cell types. Again, a larger number of miRNAs was found to be uniquely regulated, with 11 (ASC), 32 (HDF), 2 (HDMVEC), 45 (HUVEC), and 49 (RPTEC) miRNAs upregulated (FDR < 0.15) only in a single cell type. Likewise, 17 (ASC), 33 (HDF), 7 (HDMVEC), 52 (HUVEC) and 49 (RPTEC) miRNAs were downregulated in a single cell type (Figure 1d).

Heatmaps and lists of all miRNAs and mRNAs regulated uniquely in a single cell type are included in Extended Data Figure 2 and Supplementary Tables 1 to 4.

### A core miRNA-mRNA expression network is regulated across different senescent cell types

To identify a potential core miRNA-mRNA expression network of senescence, mRNAs and miRNAs that changed in at least four out of five cell types were used for correlation analysis. miRNAs upregulated (n=49) and mRNAs downregulated (n=165) and, *vice versa*, miRNAs downregulated (n=39) and mRNAs upregulated (n=240) in at least four cell types were correlated. All significant miRNA-mRNA correlation pairs (FDR < 0.01) were used for visualization of a miRNA-mRNA correlation network. 29 miRNAs (upregulated) and 28 mRNAs (downregulated) and 21 miRNAs (downregulated) and 38 mRNAs (upregulated) remained in the respective networks using this criterion. Both sets of 28 and 38 mRNAs were used for pathway enrichment analysis (Figure 1e-f).

The correlation analysis of senescent cell upregulated miRNAs and downregulated mRNAs identified two genes PATL1 (n=18) and RPS11 (n=10) which were involved in most miRNA-mRNA interactions (Figure 1e). The miRNA with the most interactions was miR-184 (n=16). The top GO terms in the category ‘Biological Process’ using the mRNAs in the network were *Cytoplasmic translation* and *Ribosomal small subunit biogenesis* (Figure 1e).

In the *vice versa* correlation analysis of senescence downregulated miRNAs and upregulated mRNAs, the two mRNAs showing the highest number of interactions with miRNAs were RRAS (n=7) and PSTPIP2 (n=4) (Figure 1f). The miRNA showing most interactions was miR-454-3p (n=11). The top GO ‘Biological Process’ term enriched using mRNAs in the network was *Pyrimidine dimer repair*, consistent with a persistent DNA damage response during senescence (Figure 1f)^2^.

### High-fat diet induced senescent-cell burden in fat tissue results in a distinct miRNA expression profile that is reversed by p21^high^ senescent-cell clearance

Next, we tested whether: 1) miRNAs regulated during senescence *in-vitro* are detected as differentially expressed *in-vivo* following increases in senescent-cell burden and 2) if these changes can be reversed by senescent-cell clearance. For this purpose, we took advantage of the recently developed *PLD* transgenic mouse model^29^ that allows Tamoxifen inducible clearance of p21 highly (p21^high^) expressing senescent cells that accumulate in fat tissue of high-fat diet (HFD) fed mice^30^.

*PLD*-mice were put on a HFD at two months of age, and monthly vehicle or tamoxifen treatments started one month after the initiation of HFD feeding. Fat tissue samples were harvested after four months of HFD feeding (Figure 2a). As previously reported, HFD feeding resulted in a significant increase in p21 and p16 expression in fat tissue that was reversed by Tamoxifen induced p21^high^ senescent-cell clearance as assessed by RT-qPCR (Figure 2b).

**Figure 2:**
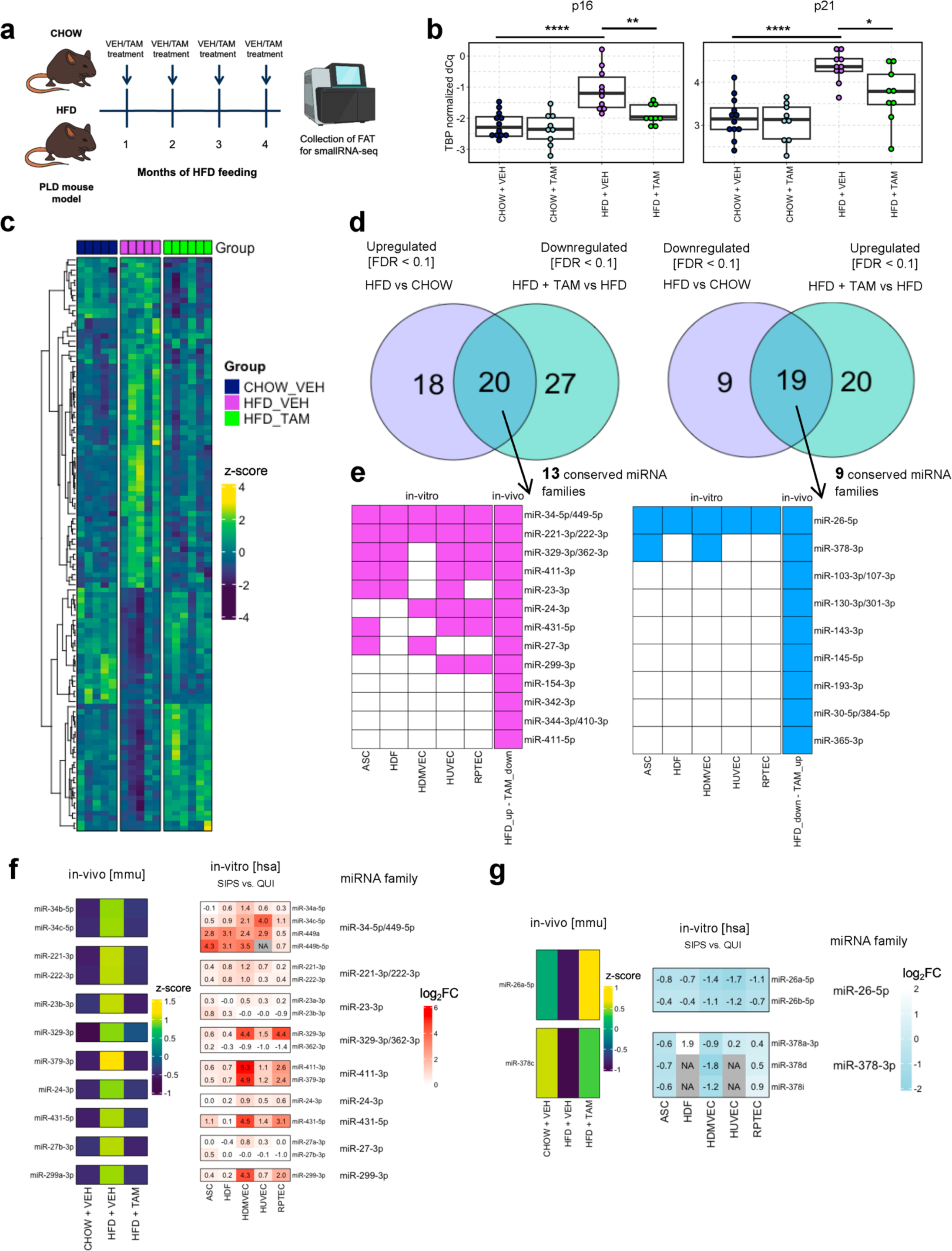
MiRNA profiles dynamically respond to changes in HFD-induced senescent-cell burden *in-vivo*. **a,** Experimental scheme for inducing and reversing senescent-cell burden in *PLD* mice by high-fat diet feeding followed by vehicle or tamoxifen injections, respectively. **b,** RT-qPCR analysis of p16 and p21 in fat tissue. **c,** Heatmap analysis using RPM values of 152 miRNAs differentially expressed in HFD VEH *vs*. CHOW VEH (FDR < 0.1) and/or HFD TAM *vs*. HFD VEH (FDR < 0.1). **d,** VENN diagram showing the overlap of miRNAs differentially regulated for the two contrasts. **e,** Comparison of conserved miRNA families up – or downregulated (FDR < 0.1), respectively, in fat tissue *in-vivo* and in different senescent cell types *in-vitro* (FDR < 0.15). **f,** Comparison of effect sizes observed for miRNAs contained in 9 miRNA families upregulated *in-vivo* and *in-vitro*. **g,** Comparison of effect sizes for miRNAs contained in 2 miRNA families downregulated *in-vivo* and *in-vitro*.

A total of 152 miRNAs was detected as differentially expressed in fat tissue following HFD feeding (HFD VEH *vs*. CHOW VEH) and Tamoxifen-induced p21^high^ senescent-cell clearance (HFD TAM *vs*. HFD VEH). Interestingly, the HFD induced changes in miRNA expression were largely reversed back to the expression pattern of chow diet fed control animals after Tamoxifen induced senescent cell clearing in HFD animals as visualized by heatmap analysis (Figure 2c).

In more detail, 66 miRNAs were significantly regulated (38 up- and 28 downregulated) following HFD feeding (HFD VEH *vs*. CHOW VEH) and 86 miRNAs regulated (39 up- and 47 downregulated) following Tamoxifen-induced p21^high^ senescent-cell clearance (FDR < 0.1). The differential regulation of 59% (n=39) out of the 66 miRNAs that changed in fat tissue following HFD was reversed by p21^high^ senescent-cell clearance (Figure 2d), reinforcing that p21^high^ senescent-cell clearance in HFD-fed mice was able to largely restore miRNA profiles to the levels observed in lean animals.

To compare the regulation of miRNAs in the *PLD* model to our *in-vitro* data, the sets of 20 & 19 miRNAs identified in the Venn analysis that were up- or downregulated by HFD and rescued by TAM, respectively (Figure 2d), were grouped into miRNA families according to miRNA seed sequences using the Targetscan database. These 20 & 19 miRNAs comprised 18 & 17 different seed families, out of which 13 & 9 seed families were conserved between mouse and human and used for subsequent comparisons (Figure 2e).

9 out of 13 conserved upregulated miRNA families were also detected as being upregulated (FDR < 0.15) in at least two different senescent cell types, while 2 out of 9 conserved downregulated miRNA families were also downregulated (FDR < 0.15) in at least two different senescent cell types (Figure 2e).

These 9 & 2 miRNA families comprised 17 & 5 human miRNAs, respectively, which were used for a more detailed side-by-side comparison. The regulation *in-vivo* in fat tissue and *in-vitro* in five human senescent-cell types was compared (Figure 2 f & g), revealing marked commonalities between the two datasets, particularly for upregulated miRNAs. This suggests that: i) the dynamic regulation of senescence-related miRNAs (*senomiRs*) can be detected in bulk tissues and ii) that *senomiRs* might be valuable surrogate biomarkers for senescence-burden *in-vivo*.

### Changes of cell-cycle- and SASP-related senescence markers between different tissues during natural aging

Next, we aimed to integrate microRNA data from stress (DOXO, *in-vitro*) and diet (HFD, *in-vivo*) induced senescence with data from aged mice, in which the senescent cell burden is known to be naturally increased^31^. Therefore, we harvested samples of seven different tissues (brain, fat, kidney, liver, lung, muscle, and skin) from 5-months-young (n=10) and 25-months-old *C57/BL6* male mice (n=10 per group).

We isolated RNA from all samples and used RT-qPCR analysis of two cell-cycle related senescence markers (p16 and p21) and three markers related to the SASP (IL-6, TNFα, MCP-1) to confirm an increased senescent-cell burden in all seven tissues

In all tissues, p16 expression was significantly upregulated. The highest inductions of p16 expression in aged (n=10) *vs*. young (n=10) animals were observed in the kidney (p < 0.0001, log_2_FC=3.5), skin (p < 0.0001, log_2_FC = 2.8), liver (p < 0.0001, log_2_FC = 2.7), and brain (p < 0.0001, log_2_FC = 1.6) followed by muscle (p < 0.01, log_2_FC = 1.8), fat tissue (p < 0.001, log_2_FC = 1.3), and lung (p < 0.05, log_2_FC = 0.7).

The expression of p21 was only upregulated in a subset of tissues, namely kidney (p < 0.0001, log_2_FC = 1.3), muscle (p < 0.001, log_2_FC = 1.0) and liver (p < 0.01, log_2_FC = 1.5). No changes in p21 expression were detected in skin (p = 0.11, log_2_FC = -0.2), lung (p = 0.83, log_2_FC = -0.04), and fat tissue (p = 0.34, log_2_FC = -0.3), while p21 showed a moderate but significant decrease with aging in brain (p < 0.05, log_2_FC = -0.3).

All tissues but muscle and skin showed at least one upregulated SASP factor mRNA. IL-6 was upregulated in kidney (p < 0.001, log_2_FC = 2.0) and brain (p < 0.05, log_2_FC = 0.6). TNFα was upregulated in liver (p < 0.001, log_2_FC = 1.9), kidney (p < 0.0001, log_2_FC = 1.6), lung (p < 0.0001, log_2_FC = 1.2), brain (p < 0.001, log_2_FC = 1.7), and fat (p < 0.01, log_2_FC = 1.1). MCP-1 was upregulated in liver (p < 0.05, log_2_FC = 1.4), kidney (p < 0.0001, log_2_FC = 2.0), lung (p < 0.01, log_2_FC = 0.6), and brain (p < 0.0001, log_2_FC = 1.3).

Based on reports that senescence can spread in a systemic manner through SASP-related signaling between tissues^32^, we wanted to take advantage of this dataset by exploring links of senescence markers among tissues. All senescence markers significantly (p < 0.05) regulated between aged and young animals were used to create a correlation matrix (Figure 3b). By visualizing all significant correlations in a network (Figure 3c), we observed that senescence markers in kidney and liver samples showed connections with all other tissues analyzed. Kidney, with 48 interactions with other tissues, ranked highest, followed by liver with 33. The single gene that showed most interactions with other senescence markers quantitated here was p16 expression in skin with 16 interactions. This suggests a link between skin p16 levels and systemic senescence burden in aged male mice. However, whether skin p16 transcript levels could indeed be a useful surrogate marker for systemic senescent cell burden needs more detailed investigation.

**Figure 3:**
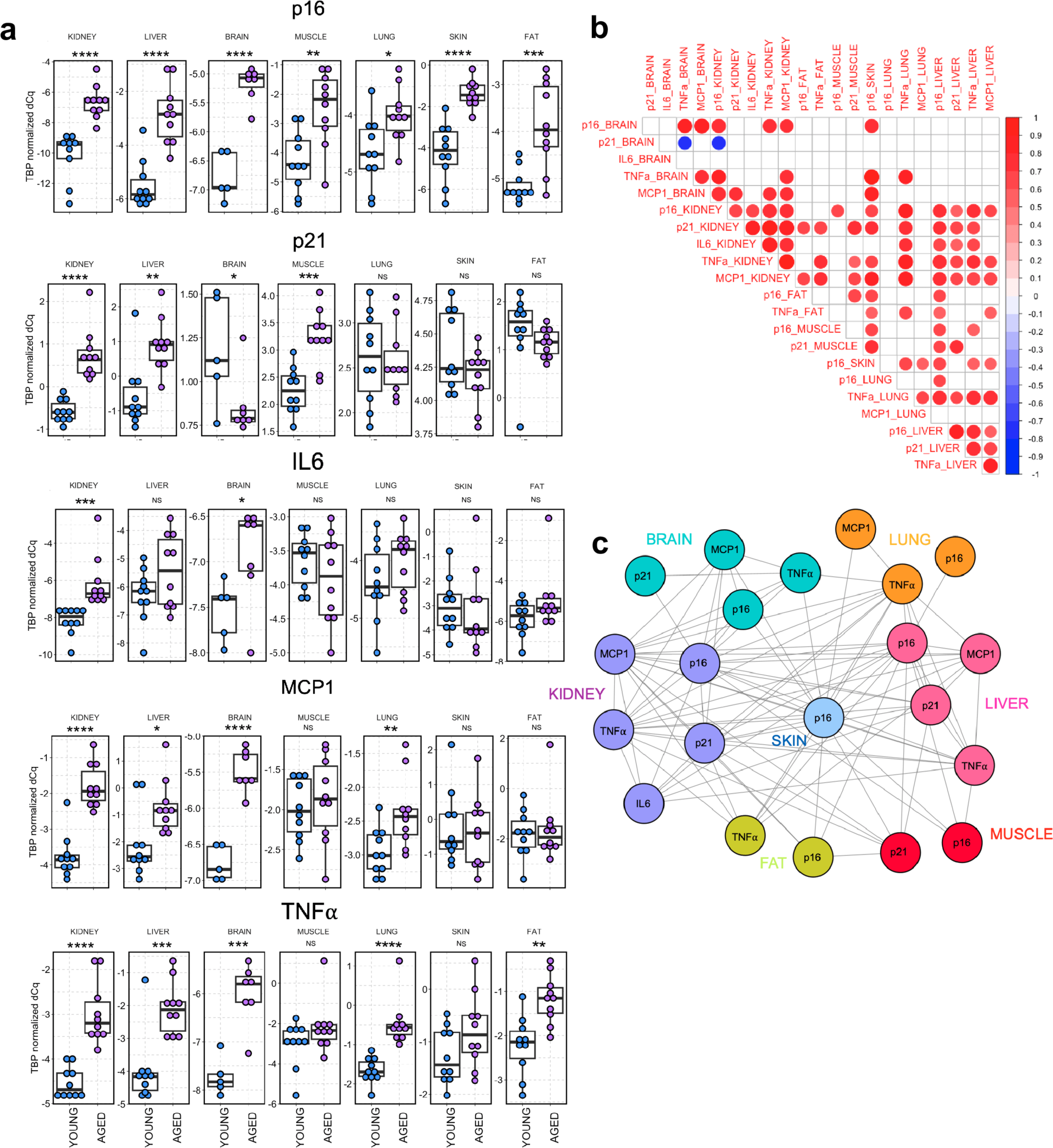
Senescence markers in seven tissues during aging. **a,** RT-qPCR analysis of p16, p21, IL6, MCP1, and TNFα in seven different tissues in aged (n=10) *vs*. young (n=10) mice. **b**, Correlation analysis of significantly regulated senescence markers in different tissues. PCCs for significant inter-tissue correlations (p < 0.05) are indicated. **c**, Correlation network using all significant correlations within and between tissues.

### Common age-related miRNA regulation identifies potential *senomiR* biomarker candidates for senescent cell burden

To comprehensively characterize miRNA changes in the 7 aged murine tissues studied here, small RNA sequencing was performed in a subset of the young (n=5) and aged (n=7) mice that were analyzed in Figure 3. Mice were selected based on availability of plasma samples sufficient to allow subsequent biofluid analysis.

Across all 84 libraries that were sequenced, 595 miRNAs were detectable with a mean raw read count > 10. Clustering of samples in a UMAP plot (Figure 4a) resulted in distinct clustering of tissues and homogenous clustering within all seven tissues and thus, all samples were included for differential expression analysis.

**Figure 4:**
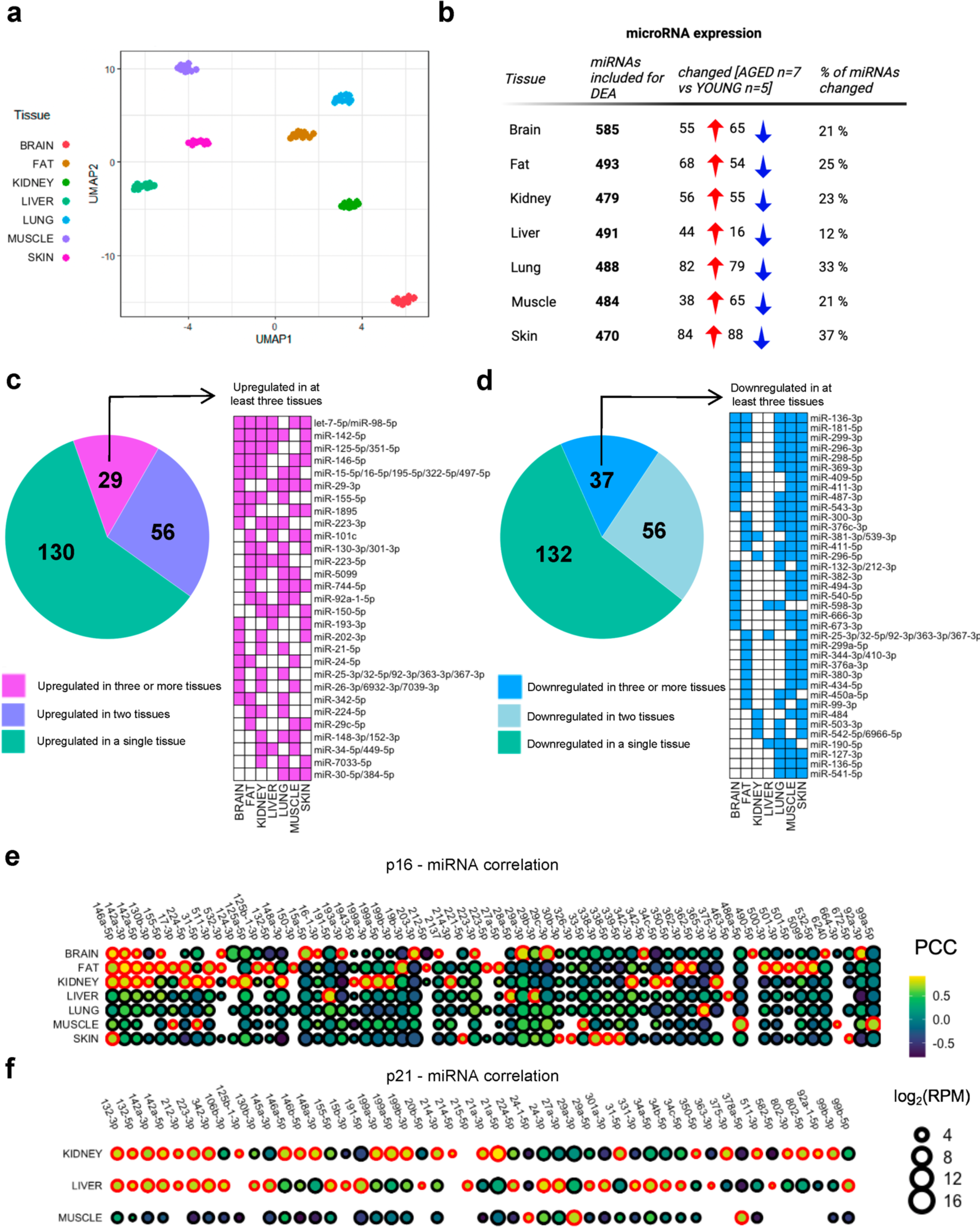
I*n-vivo* profiling of microRNA expression in seven different tissues reveals universal and tissue dependent microRNA regulation during aging. RNA was extracted from seven tissues of young (∼ 5 months, n=5) and aged (∼ 25 months, n=7) wildtype *C57BL/6J* mice and used for small RNA sequencing. **a**, UMAP plot using 592 microRNAs detectable with a mean raw read count > 10 across all 84 tissue samples included for small RNA sequencing. **b**, Differential expression analysis results for all seven tissues. The table lists the number of microRNAs that were included for DEA and the number of microRNAs significantly up- and downregulated (FDR < 0.15) in the respective tissue. **c & d**, Pie charts comparing the number of microRNA families up – or downregulated in a single tissue, two tissues, or at least three tissues. A large fraction of microRNA families is upregulated (∼ 40 %) or downregulated (∼ 41 %) in at least two different tissues during aging and a core set of 29 and 37 microRNA families is up- or downregulated, respectively, in a least three different tissues with aging. **e**, Pearson Correlation Coefficients (PCCs) for all miRNAs significantly (p < 0.01) correlated with p16 expression at least in a single tissue. **f**, PCCs for all miRNAs significantly (p < 0.01) correlated with p21 expression at least in one tissue. Significant PCCs are color coded in red.

Figure 4b lists the results of differential expression analysis. The number of miRNAs included for DEA ranged between 470 (skin) and 585 (brain) miRNAs. The number of up- or downregulated miRNAs was comparable for fat tissue (n=68 and n=54, respectively), kidney (n=56 and n=55), lung (n=82 and n=79), brain tissue (n=55 and n=65) and skin (n=84 and n=88). During liver aging, more miRNAs were upregulated (n=44) than downregulated (n=11), while the opposite was observed in muscle (n=38 and n=65, respectively) (Extended Data Figure 5a).

Considering the geroscience hypothesis of common biological processes governing age-related phenotypes, we grouped differentially regulated miRNAs according to functionally related miRNA seed families for comparisons among tissues.

A total number of 215 and 225 unique miRNA families was up- or downregulated across all seven tissues. Out of 215 upregulated miRNA families, 40% (n=85) were upregulated in at least two tissues (Figure 4c). Out of 225 downregulated miRNA families 41% (n=93) were downregulated in at least two tissues (Figure 4d). To further narrow down a core set of miRNA families with conserved regulation across multiple tissues during the aging process, all miRNA families regulated in at least three tissues were listed in Figures 4c & d. A total of 29 miRNA families was found upregulated in at least three tissues with 6 miRNA families upregulated in five out of seven tissues.

37 miRNA families were downregulated in at least three tissues. Remarkably, skin and muscle showed a large overlap of 29 out of the 37 miRNA families listed as commonly downregulated (Figure 4d).

To identify potential *senomiRs*, miRNA biomarkers of senescence, which could serve as surrogates of senescence burden, all miRNAs with a mean RPM > 10 in each tissue were correlated with p16 and p21 expression. All correlation pairs with a p-value < 0.01 were used to create dot plots for visualizing the observed correlation coefficients in each tissue (Figure 4e-f). Additional correlation plots using IL6, TNFα, and MCP1 are shown in Extended Figure 4a-c & Extended Data Figure 6.

The top five p16 *senomiRs* were miR-31-5p (PCC = 0.92, p = 2.98 x 10^-5^) and miR-148a-3p (PCC = 0.92, p = 3.00 x 10-5) in kidney samples and miR-142a-3p (PCC = 0.91, p = 3.31 x 10^-5^), miR-362-5p (PCC = 0.90, p = 8.17 x 10^-5^), and miR-142a-5p (PCC = 0.90, p = 8.57 x 10^-5^) in fat tissue. Two of these, miR-142a-3p and miR-142a-5p, were also the only *senomiRs* observed to be correlated with p16 in three tissues (brain, fat, and kidney). A single *senomiR*, miR-146a-5p, was correlated with p16 in four tissues (brain, fat, kidney, and skin). Most p16 *senomiRs* (54 out of 64) were detected as significantly correlated only in a single tissue.

The top five p21 *senomiRs* were miR-21a-5p (PCC = 0.94, p = 7.45 x 10^-6^) and miR-92a-1-5p (PCC = 0.88, p = 1.38 x 10^-4^) in kidney, miR-331-3p (PCC = 0.87, p = 2.02 x 10^-4^) and miR-24-3p (PCC = 0.86, p = 3.85 x 10^-4^) in liver, and miR-214-5p (PCC = 0.85, p = 4.21 x 10^-4^) in kidney. Seven *senomiRs* (out of 49) were correlated with p21 in both kidney and liver (miR-132-3p, mIR-132-5p, miR-142a-5p, miR-142a-3p, miR-212-3p, miR-223-3p, and miR-342-3p). Again, most p21 *senomiRs* (n=42) were correlated with p21 in a single out of the three tissues. These data support the idea that *senomiRs* might be used as biomarkers of general as well as of tissue specific senescence.

Additional comparisons of conserved miRNA families regulated in five senescent cell types *in-vitro* and seven tissues during aging *in-vivo* are included in Extended Data Figure 4d.

### senomiR levels correlate between tissues and plasma and have potential as extracellular biomarkers of cellular senescence

An important goal of this study was to identify novel extracellular biomarkers of cellular senescence using liquid biopsies. Hence, we used plasma samples collected from the same group of mice we used for tissue miRNA profiling for subsequent small RNA sequencing. Previous studies have shown that miRNA profiles recovered from total plasma and plasma derived EVs are distinct^33^. To explore the potential of either circulating miRNA fraction to reflect senescence-burden, both platelet-poor plasma (PPP) and EVs isolated from PPP using size-exclusion chromatography (SEC) from aged (n = 7) and young (n = 5) mice were used for small RNA seq.

Indeed, unsupervised PCA showed that miRNA profiles from plasma and plasma derived EVs are distinct (Figure 5a). In total plasma, we identified 45 up- and 43 down-regulated miRNAs with age. In plasma EVs 9 up- and 11 down-regulated miRNAs were identified (FDR < 0.15) (Figure 5b & Extended Data Figure 5b).

**Figure 5:**
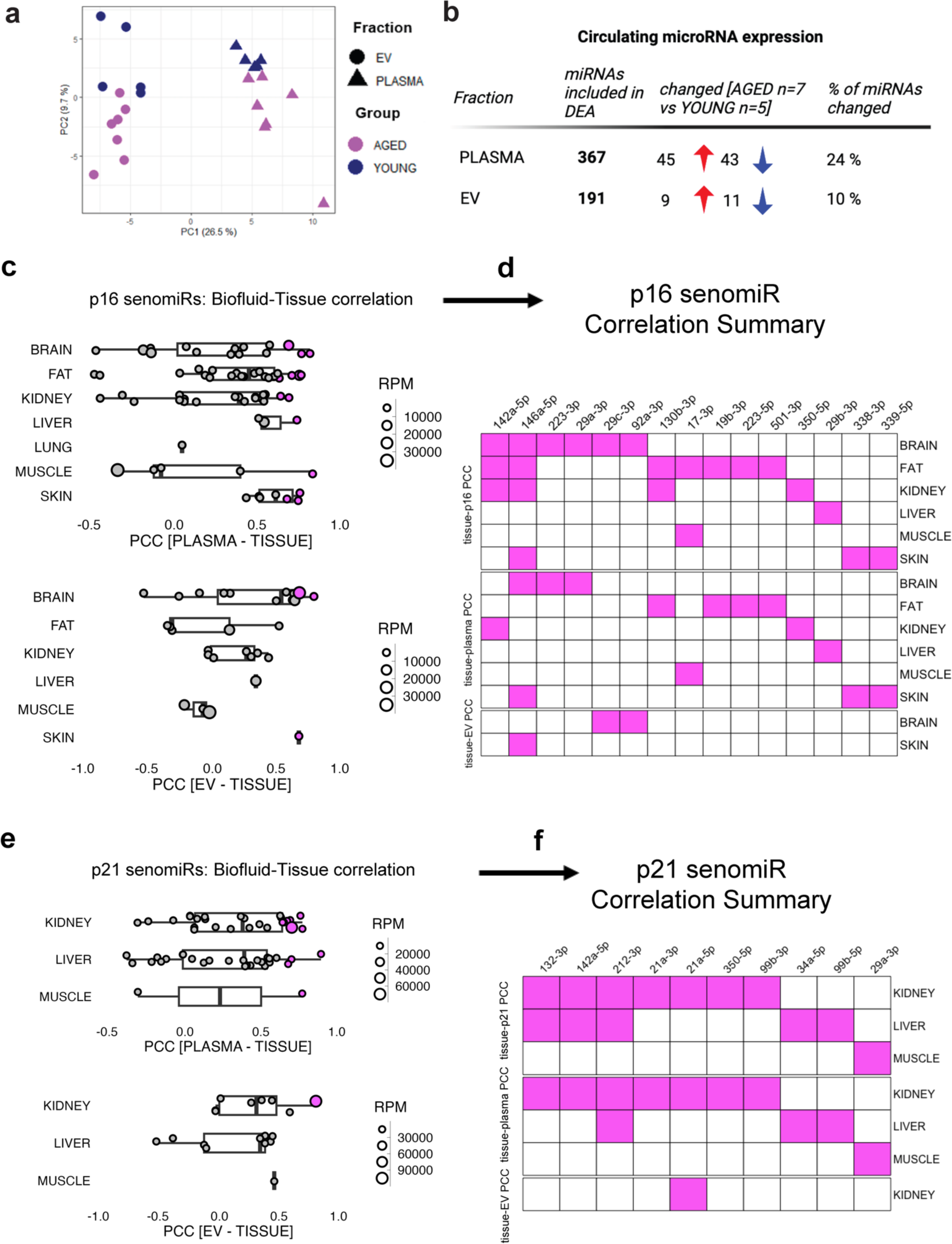
MicroRNA profiling in different fractions of the cell-free circulation of aged (∼ 25 months) and young (∼ 5 months) wildtype *C57BL/6* mice identifies potential minimally invasive microRNA biomarkers of cellular senescence. **a,** Principal Component Analysis (PCA) using plasma and plasma-derived EV miRNA expression. For unsupervised PCA only miRNAs with a mean raw read count > 100 were used. MiRNAs were ranked according to their CV% and the top 150 miRNAs were used for PCA. **b**, Differential expression results for both cell-free blood fractions analyzed. The table lists the number of miRNAs that were included for DEA and the number of miRNAs significantly up- and downregulated (FDR < 0.15) in the respective fraction. **c**, Correlation of tissue and plasma and tissue and plasma-EV miRNA levels, respectively, of p16 *senomiRs*. **d**, Summary of p16 *senomiRs* correlated with p16 in tissues and between tissues and plasma and plasma EVs, respectively. **e**, Correlation of tissue and plasma and tissue and plasma-EV miRNA levels of p21 *senomiRs*. f, Summary of p21 *senomiRs* correlated with p21 in tissues and correlated among tissues and plasma and plasma EVs, respectively. Tissue-biofluid correlations reaching a p value < 0.05 are highlighted in purple in panels c-f.

To profile circulating *senomiR* candidates that could reflect senescence burden in a specific tissue, all *senomiRs* that were significantly correlated with p16 and/or p21 on tissue level were correlated with the abundance in plasma and EVs.

Out of p16 *senomiRs*, the top tissue-plasma correlations we observed were for miR-146a-5p between plasma and brain (PCC = 0.76, p < 0.01) and skin (PCC = 0.70, p < 0.01) and for miR-17-3p between plasma and muscle (PCC = 0.78, p < 0.01). EV levels of miR-29c-3p were correlated with tissue levels in brain (PCC = 0.72, p < 0.01) (Figure 5c).

For p21 *senomiRs*, the top miRNA was miR-34a-5p in plasma that was significantly correlated with levels in liver (PCC = 0.84, p < 0.01), followed by plasma miR-132-3p (PCC = 0.72, p< 0.01) correlation with kidney and plasma miR-29a-3p (PCC = 0.72, p < 0.01) correlation with muscle. EV levels of miR-21a-5p were significantly correlated with tissue levels in kidney (PCC = 0.74, p < 0.01) (Figure 5e).

In total, 15 and 10 circulating p16 and p21 *senomiR* candidates were identified respectively and summarized in Figure 5d & Figure 5f.

## Discussion

Here, we present a comprehensive resource cataloging miRNA expression during senescence and aging using five human senescent cell types and two *in-vivo* models with increased senescence burden. We show that miRNA profiles follow senescence burden dynamically in *PLD* mice, using HFD-induced senescence and conditional p21^high^ senescent cell ablation. Moreover, we profiled miRNAs in seven different tissues and two fractions of the circulation during natural aging, exploring potential *senomiR* biomarker candidates.

We have previously shown miRNAs of the miR-17-92 cluster to be downregulated in common during senescence and aging^20^, and several other studies have reported differential miRNA regulation during cellular senescence^20,25,34–37^. However, none have systematically compared miRNA & mRNA expression profiles across multiple primary human senescent cell types.

To characterize miRNA and mRNA expression changes during senescence, we collected RNA from five different human primary cell types three weeks after Doxorubicin induced senescence. We discovered that miRNA expression is highly senescent-cell type specific, with most regulated miRNAs being unique for each cell type.

Our results from correlating miRNA and mRNA levels suggest the involvement of miRNAs in various aspects of the senescent phenotype. Commonly upregulated miRNAs such as miR-215-5p and miR-184 showed significant correlations with members of ribosomal subunits, a group of mRNAs downregulated in all five senescent cell types. This downregulation paralleled a study showing senescence-related defects in ribosomal biogenesis reinforce cell-cycle arrest^38^, raising the possibility of a connection to these miRNAs.

Regarding downregulated miRNAs and upregulated mRNAs, we observed an enrichment of GO terms related to DNA damage. Particularly miRNAs of the miR-17-92 cluster (miR-17-5p, miR-20a-5p and miR-92a-3p) exhibited correlations with genes linked to these terms. Notably, downregulation of this cluster was consistent across all five senescent cell types as previously reported by us^20^, but was only observed in fat and liver during natural aging *in-vivo*, creating an interesting connection to a report on compromised adipose tissue homeostasis in mice following knock-out of miR-17-92 family members^39^.

To ascertain the contribution of miRNAs to different phenotypic aspects of senescence, more detailed experiments using knock-out of individual miRNAs will be needed. Given that a general loss of miRNAs *via* knock-out of Dicer was shown to sensitize cells to undergo senescence^40^, it is unlikely that the miRNA pathway is essential for a general senescence response. Based on the observed senescent cell-type specificity of miRNA expression, we hypothesize it is more plausible that they play a role in efficiently orchestrating various phenotypic features such as cell-cycle arrest or cell-type specificity of the SASP^41^.

The *in-vivo* relationship of miRNA expression and senescence-burden remains underexplored. To consider miRNAs as circulating biomarkers of senescence, it is crucial to demonstrate the dynamic response of miRNA changes in bulk tissues following senescent-cell induction and clearance. We leveraged the recently developed *PLD* mouse model for this purpose, allowing for Tamoxifen inducible clearance of p21^high^ senescent cells^29^. Clearing p21^high^ senescent cells in fat tissue resulted in an almost complete rescue of miRNAs differentially expressed during HFD feeding, underscoring the broad-reaching effects of depleting this subpopulation of cells. This observation not only affirms the potential of miRNAs as surrogates of senescent-cell abundance, but also insinuates a novel functional perspective for conditions associated with increased senescence.

Furthermore, our findings in fat tissue aligned closely with miRNAs upregulated in multiple senescent cell types. The 20 mature miRNAs upregulated following HFD feeding and subsequently reversed by p21^high^ cell ablation belonged to 13 conserved miRNA seed families. Of these, 9 were also upregulated in at least two out of five senescent cell types. In this comparison, at least one member of the miR-34-5p/449-5p family was upregulated in all five senescent cell types. The relationship between this miRNA-family and expression of p21 was consistently observed in the different experimental systems we used: in aged mice, we only found the upregulation of members of the miR-34-5p/449-5p family in tissues that also showed an upregulation of p21 with age (kidney, liver, and muscle). Considering the established association of this miRNA family with the p53 pathway^42^, our observations suggest its potential as biomarker in the context of p21-dependent senescence.

Our analysis of miRNA changes in seven different tissues during natural aging was focused on i) identifying common miRNA families regulated during aging and cellular senescence and ii) mining for potential *senomiRs*, miRNA biomarkers of senescence that could be used to monitor the senescent-cell load of patients non-invasively.

The consistency of detection of *senomiR* candidates belonging to the miR-146-5p and miR-29-3p families in both analyses is noteworthy.

The levels of miR-146a-5p were correlated with p16 in multiple tissues and the abundance in the circulation (both plasma and plasma EVs) was correlated with levels in the skin. This could be of relevance, as p16 levels in skin were correlated to senescence markers in all other tissues and point to a potential role as a read-out of systemic senescence burden. We have found miR-146a-5p upregulated in HDFs, HUVECs, and RPTECs during DOXO induced senescence, in line with previous reports about increased levels of miR-146a-5p during replicative senescence^43^ and in senescent IMR90 cells^35^. Moreover, *in-vivo* overexpression of miR-146a-5p was shown to induce senescence in adipose tissue^44^ and this miRNA was also observed by others to be upregulated in the circulation during aging in humans^45^ and mice, which was prevented by putting mice on a calorie restriction diet^46^, an intervention known to decrease senescence markers in multiple tissues^31^ and in humans^47^.

Plasma levels of miR-29b-3p correlated with levels in liver, miR-29a-3p and miR-29c-3p correlated with brain levels. The miR-29 family, consisting of miR-29a, miR-29b, and miR-29c, has been shown to induce senescence in muscle, both *in-vitro* and *in-vivo*^36^. Our study revealed a remarkably widespread upregulation of miR-29-3p family members with aging in five out of seven tissues and three out of five human senescent cell types. This association with aging seems to be highly conserved across species as miR-29 was shown to be upregulated in the brain of fish^48^, mice^49^, as well as humans and macaques^50^. Given the many links between miR-29, senescence, and aging, miR-29 family members could possibly represent a valuable minimally invasive read-out of organismal senescence burden.

In summary, our study presents a valuable resource for the scientific community and sets the stage for deeper exploration of miRNAs as specific biomarkers of senescence in different tissues. We have identified a list of 15 and 10 circulating *senomiRs* potentially associated with p16 and/or p21, respectively. Notably, only three circulating *senomiRs* (miR-142a-5p, miR-350-5p, miR-29a-3p) overlapped in their association with both p16 and p21. This raises intriguing questions about the potential of miRNAs in delineating p16^high^ and p21^high^ senescent cell populations, a hypothesis that warrants detailed investigation in future studies.

## Methods

### Cell Culture

Human dermal fibroblasts (HDF), adipose-dervied stem cells (ASC), human dermal microvascular endothelial cells (HDMVEC), human umbilical vein endothelial cells (HUVEC) and human renal proximal tubule epithelial cells (RPTEC) from three healthy donors respectively were provided by Evercyte GmbH. Written informed consent was obtained from all patients before tissues samples were taken. The study was conducted in accordance with the principles of the Declaration of Helsinki.

HDFs were cultivated in DMEM (Gibco, MA, USA) supplemented with 10 % fetal bovine serum (FBS) and 4 mM L-Glutamine (Gibco, MA, USA). ASCs were cultivated in EGMTM-2 Endothelial Cell Growth Medium (Lonza, Switzerland) supplemented with 2 % FBS. HDMVECs were cultivated using the Endopan MV kit (PAN Biotech, Germany) and for HUVECs, EGMTM Endothelial Cell Growth Medium BulletKitTM (Lonza, Switzerland) supplemented with 10 % FBS was used. RPTECs were cultivated in ProxUp2 medium (Evercyte GmbH, Austria) without G418 supplement.

Cells were seeded at a density of 3500 cells/cm^2^ (day 0). For induction of senescence, cells were treated with 100 nM Doxorubicin on the day after seeding (day 1), medium was changed with fresh 100 nM Doxorubicin at day 4 until recovery was initiated on day 11 by changing to normal growth medium. Cells grown into quiescence were used as controls and were seeded on the same day as cells that were exposed to Doxorubicin. For RPTECs, the protocol was adapated to a seeding density of 20000 cells/cm2 (day 0) and treatment with 200 nM Doxorubicin.

### Senescence-associated Beta-galacotosidase staining

Senescence-associated β-galactosidase staining was performed according to a previously reported protocol^51^. Cells were stained on day 21 after seeding and imaged at a 100x magnification.

### BrdU assays

For verification of the senescence-associated growth arrest, cells were incubated for 24 hours with 10 μM BrdU (Sigma Aldrich, MO, USA). Cells were fixed in ice-cold 70 % ethanol for at least one hour at 4 °C and stored at -20 °C for subsequent antibody staining.

Cells were permeabilized with 2 M HCl and 1 % Triton X-100 (Sigma Aldrich, MO, USA), followed by neutralization with 0.1 M Na-Borat (pH 8.5). Cells were stained with a 1:50 dilution of anti-BrdU antibody (BD Biosciences, USA) in TBS and incubated for 30 minutes at room temperature. Next, cells were stained with a 1:100 dilution of anti-mouse FITC-conjugated antibody (Sigma Aldrich, MO, USA). Finally, cells were stained with DAPI and used for microscopic imaging of BrdU and DAPI signals.

### Animals

For the aging study at the Mayo Clinic Rochester (USA), all animal experiments were performed according to protocols approved by the Institutional Animal Care and Use Committee (IACUC). Wild type *C57BL/6* mice were obtained from the National Institute on Aging (NIA) and all mice were maintained in a pathogen free facility at 23 – 24 °C under a 12h light, 12h dark regimen with free access to a NCD (standard mouse diet with 20 % protein, 5 % fat, and 6 % fiber, Lab diet 5053, St. Louis, MO) and water. All mice were housed at 2 to 5 mice per static autoclaved HEPA ventilated microisolator cage (27 x 16.5 x 15.5 cm) with autoclaved Enrich-o’Cobs (The Anderson Incorporated) for bedding.

For the high-fat diet experiment, mice were a fed a diet of the following caloric composition: 60 kcal% fat, 20 kcal% carbohydrate, and 20 kcal% protein (Research Diets, New Brunswick, NJ). Mice received four months of high-fat diet feeding (starting at two months of age). For Tamoxifen treatments, mice received 2 mg tamoxifen dissolved in corn oil (Sigma-Aldrich, St. Louis, MO), vehicle treated mice received equal volumes of corn oil.

### RNA isolation from tissue samples

Tissue samples were minced using a razor blade in a petri dish placed on a block of dry ice to prevent thawing. Tissue pieces were added to 1.5 ml tubes filled with ceramic beads, filled with 1000 μL of Qiazol, and homogenized by shaking for 1 minute. After homogenization, the samples were centrifuged for 30 seconds at 8000 x g and 700 μL of tissue lysate were transferred to a new tube. 140 μL of Chloroform were subsequently added, followed by cooled centrifugation at 12,000 x g for 15 minutes at 4°C. 300 μL of the upper aqueous phase were transferred to a new tube and mixed with 450 μL of ethanol. Samples were transferred to a miRNeasy mini column and washed with RPE and RWT buffer. Finally, total RNA was eluted in 30 μL nuclease free water and stored at -80°C to await further analysis.

### mRNA RT-qPCR analysis

For gene expression analysis in tissues of aged mice, RNA was reverse transcribed using a M-MLV reverse transcriptase kit (Thermo Fisher Scientific) following the manufacturer’s instructions. The following amounts of RNA were used for the cDNA reactions: 100 ng (muscle, fat tissue, and lung), 200 ng (skin), 400 ng (kidney, liver, and brain). TaqMan fast advanced master mix (Thermo Fisher Scientific) was used for real-time PCR. The following primers were used and purchased from Thermo Fisher Scientific: TATA-binding protein (TBP) (Mm01277042_ m1), CDKN2A (p16^INK4a^) (Mm00494449_m1) and CDKN1A (p21^Cip1^) (Mm04205640_g1). PCR amplification was performed in a 384 well format in a QuantStudio 7 instrument (Applied Biosystems) at the following settings: 95 °C for 20 s followed by 40 cycles of 95 °C for 1 s and 60 °C for 20 s. Cq-values were calculated using default software settings for baseline and threshold determination.

### Statistical analysis

mRNA sequencing data were processed as recommended by Lexogen. FastQC v0.11.9 and multiQC v.10 were used to check the quality of the demultiplexed raw as well as the preprocessed data. Cutadapt v.3.3 was used to trim the RealSeq adapter sequence from smallRNA sequencing derived reads and further remove reads with a quality score below 30 and shorter than 17 nucleotides. For mRNA sequencing data trimming, and filtering were conducted using bbmap and the function bbduk v.38.90, the same cut-offs as for small RNA preprocessing were used. Small RNA sequencing data were mapped against the human genomic reference GRCh38.p12 provided by Ensembl using bowtie v1.3.0. All mapping reads were further processed by miRDeep2 v.2.0.1.2 by mapping against miRBase v22.1. allowing for one mismatch and, depending on the experiment, filtering for human (hsa) or murine (mmu) specific microRNAs. All genomic non microRNA related reads were investigated for general RNA composition using bowtie v.1.3.0 and RNAcentral. Sequencing data of mRNA were first mapped against the human genomic reference GRCh38.p12 provided by Ensembl using STAR v.2.7.7a. Feature counts were generated using HTSeqCount v.0.12.4. Downstream analysis of annotated reads was performed using R v4.04 and packages derived from CRAN or Bioconductor.

Differential expression analysis was performed using the package edgeR and the independent filtering method of DESeq2. The DESeq2 filtering method was adapted in order to remove low abundant miRNAs. Raw count data and a data frame containing group information were passed to the Bioconductor package edgeR. The design matrix was prepared to compare two groups.

For all other univariate statistical analyses, GraphPad Prism version 8.0.0 (GraphPad Software, San Diego, California USA, www.graphpad.com) was used. As indicated either unpaired t-tests or 1-way ANOVA in conjunction with Tukey’s *post-hoc* test were used for assessing statistical significance. Data were visualized using the R packages *ggplot2*, *ComplexHeatmap*, and *venn*. Comparisons of regulated features between conditions were performed using the R package *upsetR*. For the analyses on miRNA families, data on miRNA seed families were downloaded from https://www.targetscan.org/mmu_80/.

### microRNA-mRNA network construction

To assess potential microRNA-mRNA interactions in different senescent cell types, microRNAs upregulated in at least four out of five senescent cell types were correlated with genes downregulated in at least four out of five senescent cell types and *vice versa* in all 30 *in-vitro* samples generated. All microRNA-mRNA correlations reaching a threshold of an adjusted p-value < 0.05 were used to generate a network using Cytoscape 3.8.1.^52^. All mRNAs that remained in the networks were used for GO term enrichment in the category “Biological Process” using ShinyGO (v0.80)^53^.

### NGS library preparation

For small RNA library preparations, the RealSeq-Biofluids library kit for Illumina sequencing (RealSeq Biosciences, CA, USA) was used according to the recommendations by the manufacturer. For tissue and cell samples, 100 ng total RNA diluted in 8.5 μl plus 1.0 μl of miND® spike-ins^54,55^ were used as starting material. Adapter-ligated libraries were circularized, reverse transcribed, and amplified. PCR amplification with Illumina primers of libraries was performed using 23 cycles. In total, 30 libraries were generated using *in-vitro* samples from RNA isolated from cells. 100 libraries were generated from murine tissue samples and libraries were quantified using the Agilent DNA 1000 kit (Agilent Technologies, USA).

The generated libraries were diluted and pooled to equimolar concentrations and size-selected with the BluePippin system using a 3 % agarose cassette (Sage Science, USA) to remove adapter dimers and other contaminating DNA fragments outside the target range. The pooled and purified libraries were quantified using the Agilent High Sensitivity DNA kit (Agilent Technologies, USA). The pool of *in-vitro* cellular RNA libraries was sequenced on an Illumina NextSeq550 (single-read, 75 bp) instrument. The sequencing run was performed according to the manufacturer’s protocol at the Vienna BioCenter Next Generation Sequencing core facility in Vienna, Austria.

The 84 tissue libraries generated for the aging study were separated into two pools and were sequenced on two Illumina NovaSeq6000 lanes (SPrime, single-read, 100 bp) according to the manufacturer’s protocol at the University of Minnesota Genomics Center (UMGC), Minneapolis, USA. The 16 fat tissue libraries generated from *PLD* mice were sequenced on a NextSeq550 (single-read, 75 bp) instrument according to the manufacturer’s instructions at the UMGC, Minneapolis, USA.

Mapping statistics for all sequencing runs are shown in Supplementary Figure 1.

### mRNA library preparation

The QuantSeq 3’ mRNA-Seq library kit FWD for Illumina (Lexogen, Austria) was used to generate mRNA libraries using 100 ng of total RNA as input material. 17 cycles were used for PCR amplification of adapter-ligated libraries. Library concentrations were quantified using the Agilent High Sensitivity DNA kit (Agilent Technologies, USA). An equimolar pool consisting of 24 samples was prepared and sequenced on an NextSeq550 (single-read, 75 bp).

## Acknowledgements

We would like to acknowledge our funding sources: Austrian Science Fund (Österreichischer Wissenschaftsfonds; FWF) Grant DOIs: 10.55776/P35268 (J.G.); 10.55776/P35382 (M.O.), 10.55776/P36483 (M.O) 10.55776/P37321 (M.O.), a Federation of European Biochemical Societies (FEBS) Excellence Award (M.O.), a EUREKA-Eurostars Grant E!828 AB-SENS (M.H.) and National Institute on Aging grant R37AG013925 (J.L.K.), the Connor Fund (J.L.K.), Robert J. and Theresa W. Ryan (J.L.K.), and the Noaber Foundation (J.L.K.).

## Author contributions

J.G., M.H., Mo.W., J.L.K., and T.T. conceived and designed the study and revised the manuscript. Mo.W. performed most experiments, analyzed and interpreted all data and wrote the original draft of the manuscript. Ma.W., T.K., M.P., and B.M. contributed to *in-vitro* data collection. S.C., C.L.I., T.P., K.O.J., N.G., A.X., M.S. and L.W. supported *in-vivo* experiments. M.X. shared *PLD* mice and advised on related experiments. J.G., M.H., J.L.K., T.T., M.P., A.D., K.K., I.L., R.G., and M.O. analyzed and interpreted the results. All authors reviewed the results and approved the final version of the manuscript.

## Corresponding author

Correspondence to Johannes Grillari.

## Extended Data Figures

**Extended Data Figure 1:**
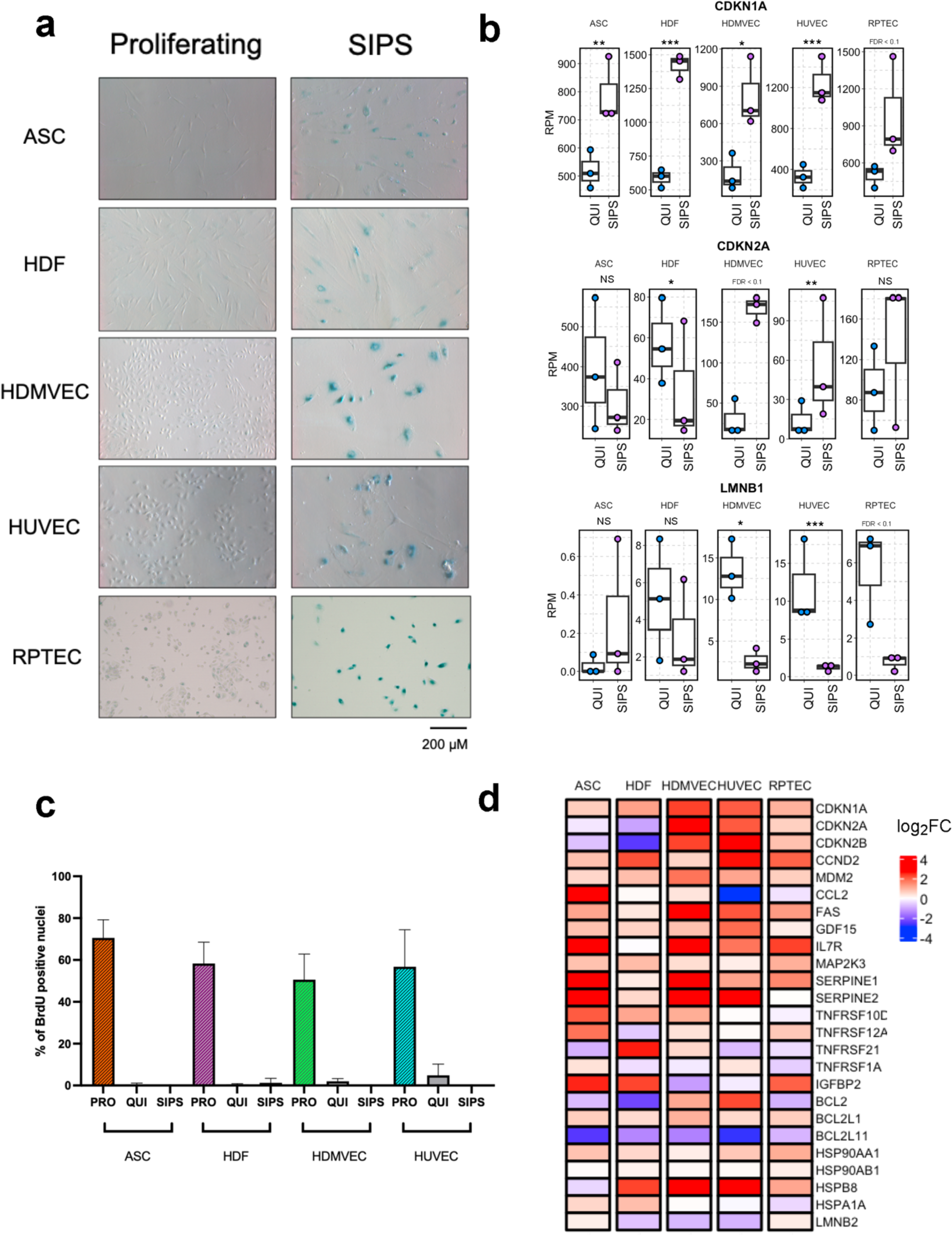
**a**, Senescence-associated Beta-Galactosidase staining in all five cell types **b**, Expression of selected senescence markers CDKN1A, CDKN2A, and LMNB1 in all five cell types **c**, BrdU incorporation assay in proliferating (PRO), quiescent (QUI) and stress- induced senescent (SIPS) cells **d**, log_2_- fold changes for panel of senescence markers.

**Extended Data Figure 2:**
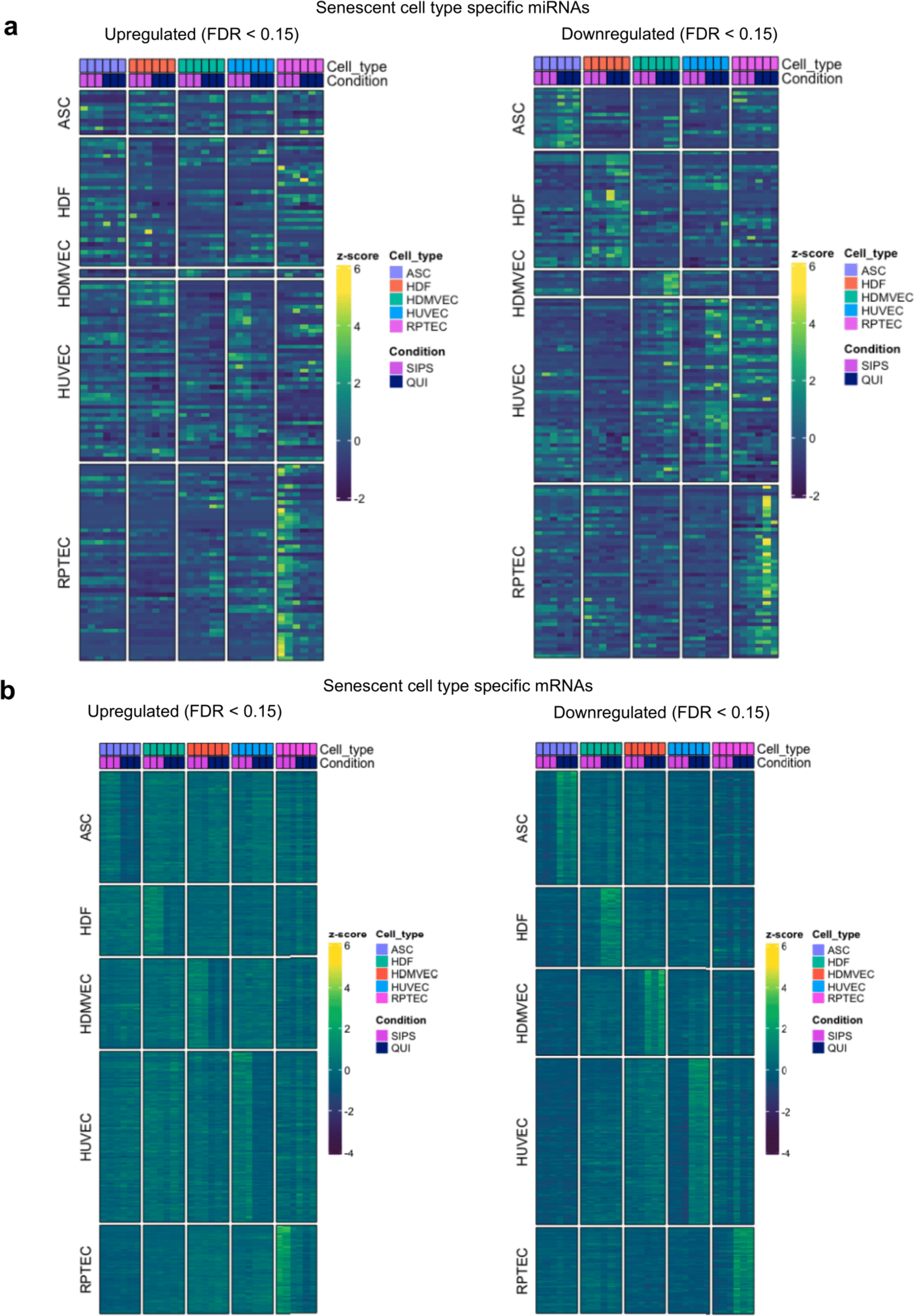
**a,** Heatmaps on uniquely up- and downregulated (FDR < 0.15) miRNAs in individual senescent cell types. **b,** Heatmaps on uniquely up- or downregulated (FDR < 0.15) mRNAs in individual senescent cell types. For all heatmaps, row annotations specify the senescent cell type of the respective uniquely regulated miRNA/mRNA set.

**Extended Data Figure 3:**
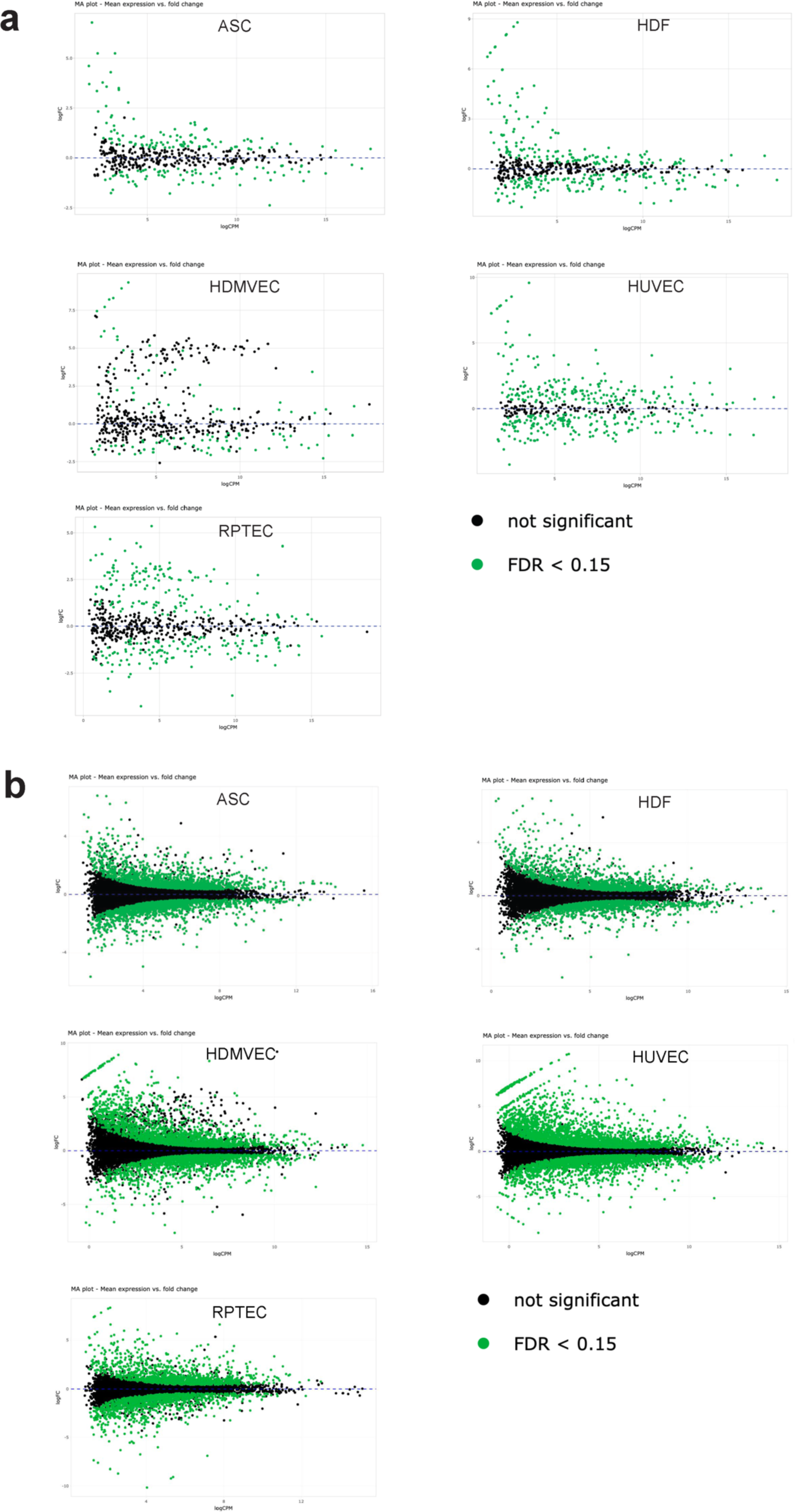
**a,** MA plots of miRNA expression in all five cell types. **b,** MA plots of mRNA expression in all five senescent cell types. Differentially expressed (FDR < 0.15) miRNAs and mRNAs respecrtively are highlighted in green.

**Extended Data Figure 4:**
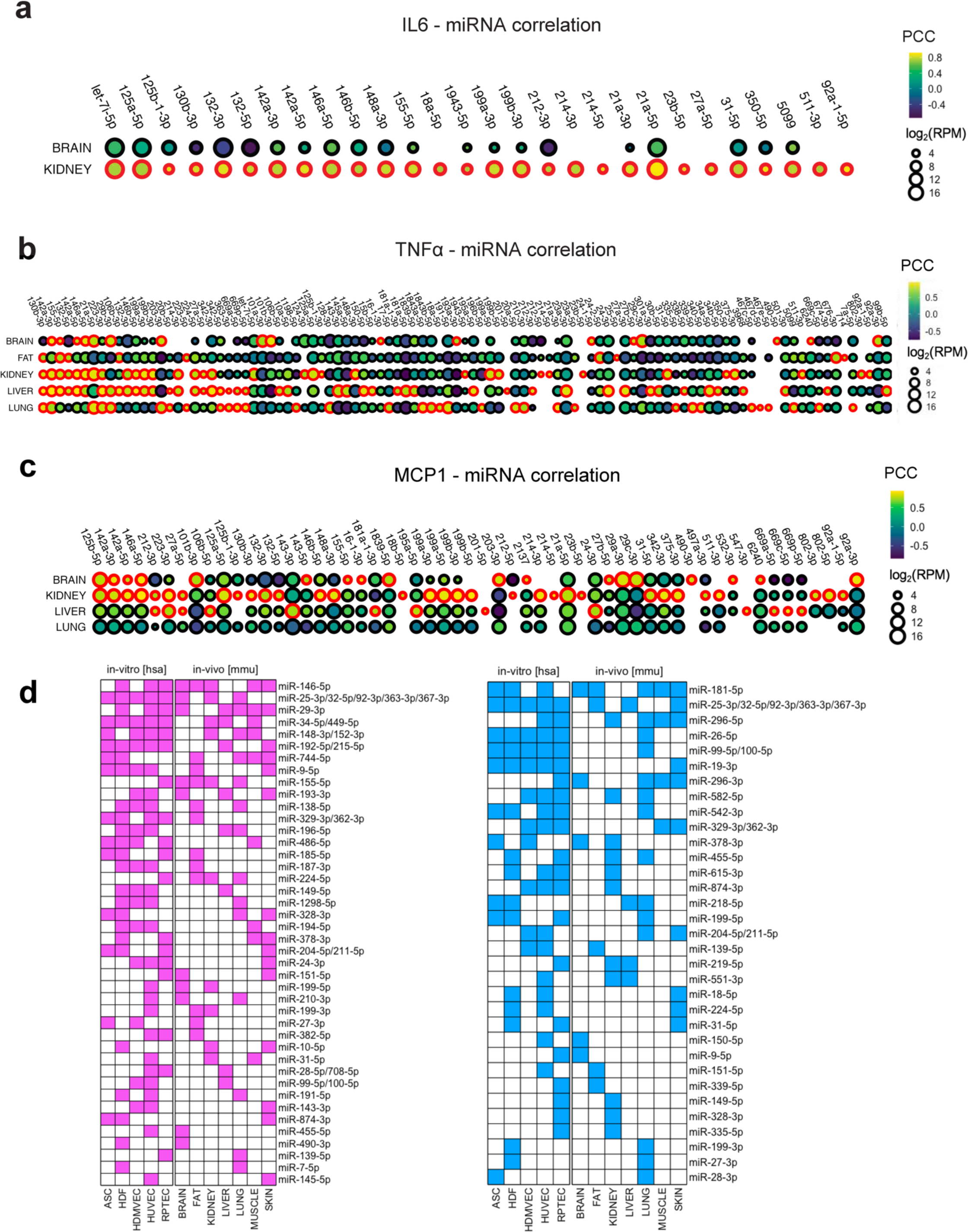
Pearson Correlation Coefficients (PCCs) for all miRNAs significantly (p < 0.01) correlated with **a,** IL6 **b,** TNFα and **c,** MCP-1 in at least a single tissue. Comparison of conserved **d,** upregulated and **e,** downregulated miRNA families in five human senescent cell types *in-vitro* (FDR < 0.15) and seven different mouse tissues *in-vivo* (FDR<0.15).

**Extended Data Figure 5:**
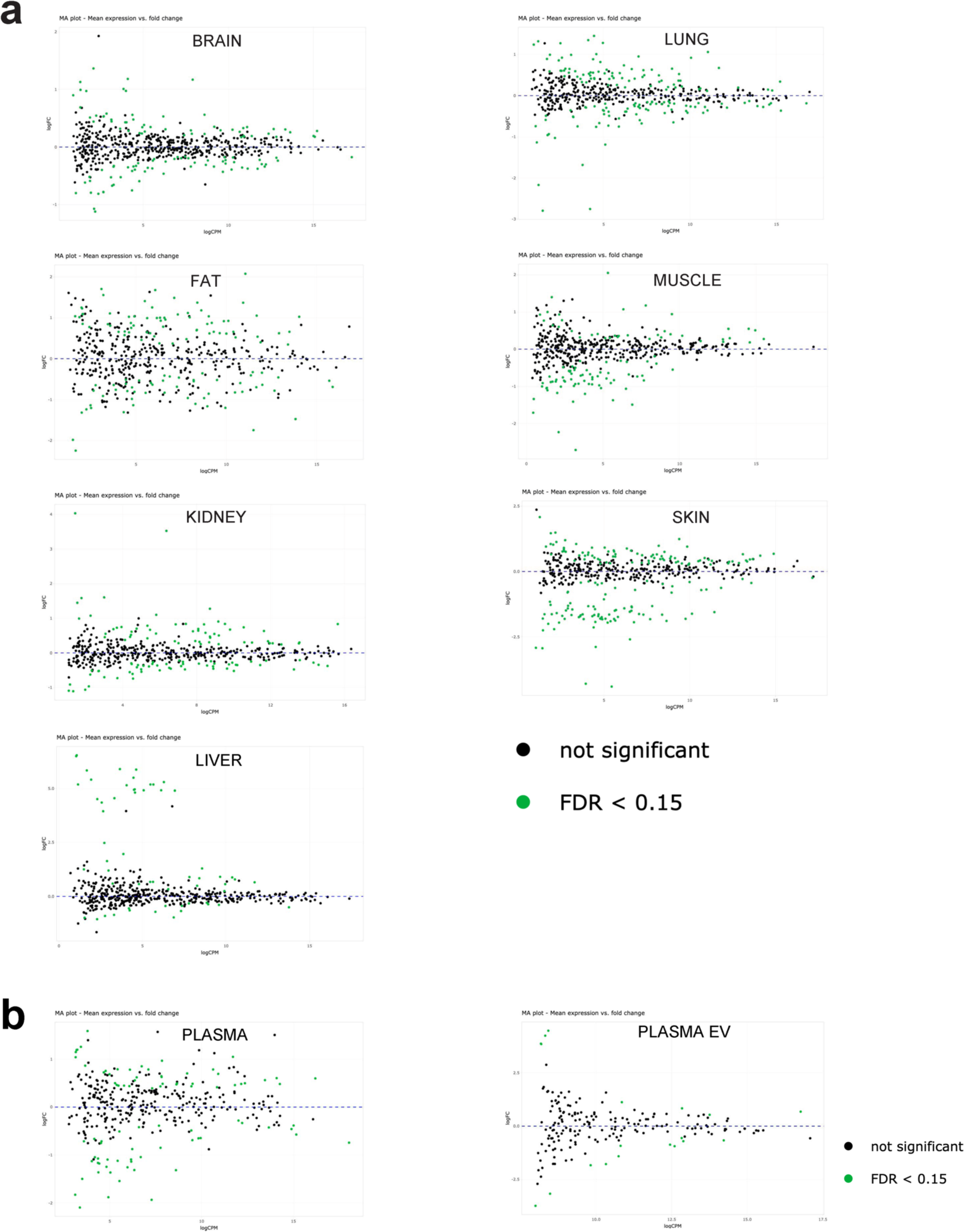
MA plots of miRNA expression in **a,** all seven tissues analyzed in the aging study and **b,** plasma and plasma derived EVs included in the aging study. Differentially expressed (FDR < 0.15) miRNAs are highlighted in green.

**Extended Data Figure 6:**
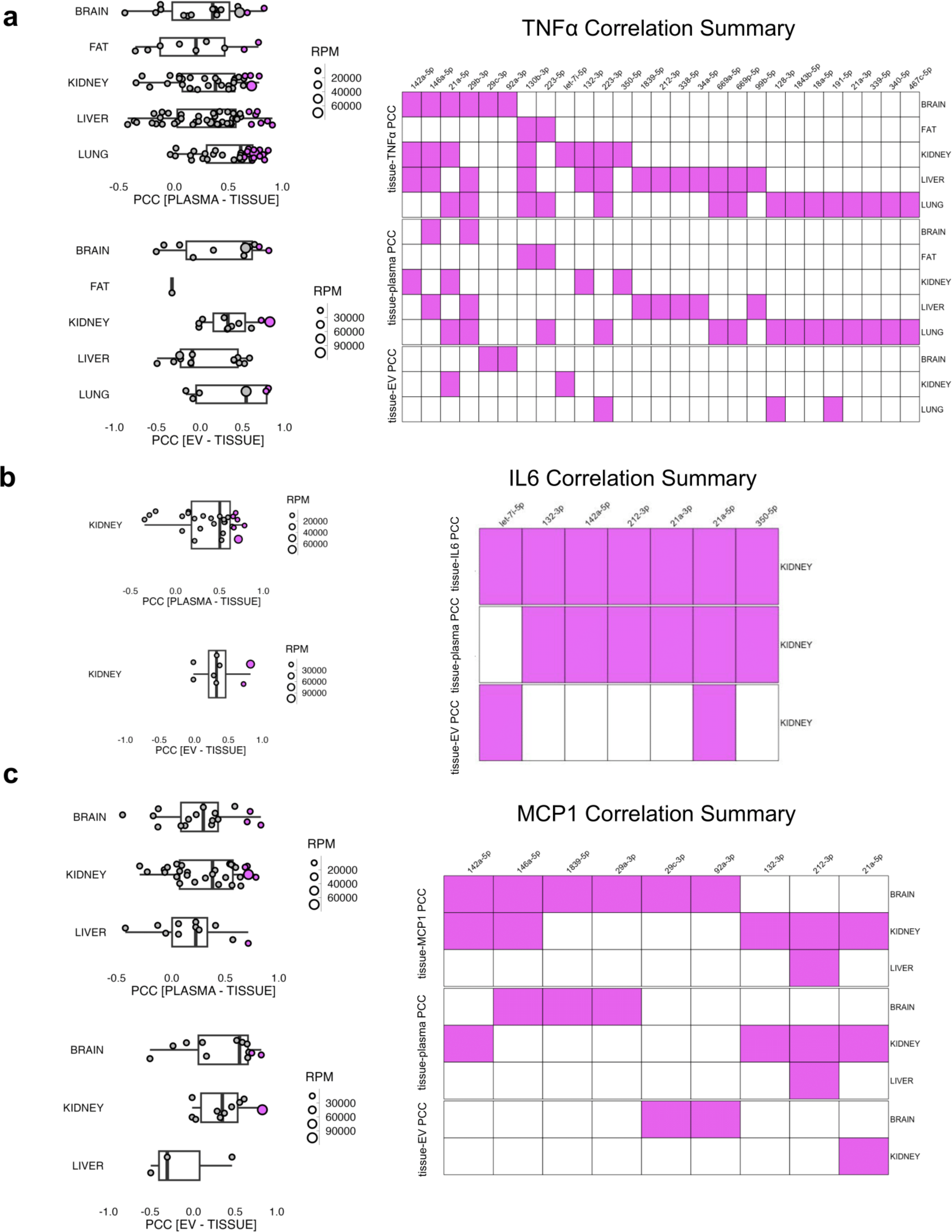
Correlation of tissue and plasma and tissue and plasma-EV miRNA levels, respectively, of **a,** IL6-correlated miRNAs **b,** TNFα-correlated miRNAs, and **c,** MCP-1 correlated miRNAs. Tissue-biofluid correlations reaching a p-value < 0.05 are highlighted in purple in panels a-c.

## Supplementary Figures

**Supplementary Figure 1:**
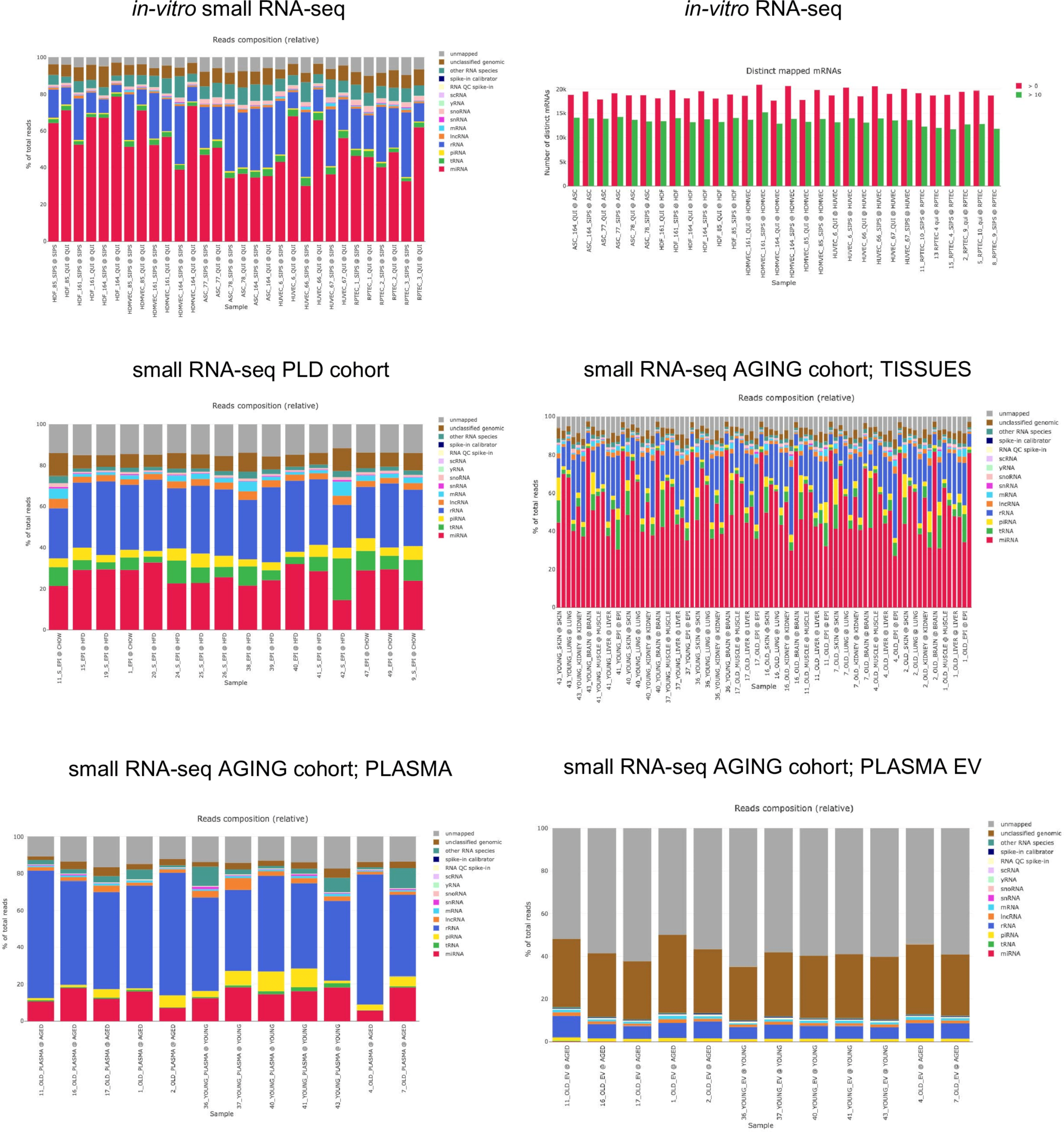
Mapping statistics for all sequencing runs.

## Supplementary Tables

**Supplementary Table 1:** List of all mRNAs that were detected as upregulated (FDR < 0.15) in each cell type. Table refers to the entire list of upregulated mRNAs (1 = ‘upregulated’, 0 = ‘not upregulated’ for each cell type).

**Supplementary Table 2**: List of all mRNAs that were detected as downregulated (FDR < 0.15) in each cell type. Table refers to the entire list of upregulated mRNAs (1 = ‘downregulated’, 0 = ‘not downregulated’ for each cell type).

**Supplementary Table 3**: List of all miRNAs that were detected as upregulated (FDR < 0.15) in each cell type. Table refers to the entire list of upregulated miRNAs (1 = ‘upregulated’, 0 = ‘not upregulated’ for each cell type).

**Supplementary Table 4**: List of all miRNAs that were detected as downregulated (FDR < 0.15) in each cell type. Table refers to the entire list of upregulated miRNAs (1 = ‘downregulated’, 0 = ‘not downregulated’ for each cell type).

## References

1. Chaib, S., Tchkonia, T. & Kirkland, J. L. Cellular senescence and senolytics: the path to the clinic. Nat Med 28, 1556–1568 (2022).

2. Zhu, Y. et al. Past and Future Directions for Research on Cellular Senescence. Cold Spring Harb Perspect Med 14, a041205 (2024).

3. Lushchak, O., Schosserer, M. & Grillari, J. Senopathies—Diseases Associated with Cellular Senescence. Biomolecules 13, 966 (2023).

4. Zhu, Y. et al. The Achilles’ heel of senescent cells: from transcriptome to senolytic drugs. Aging Cell 14, 644–658 (2015).

5. Zhu, Y. et al. Identification of a novel senolytic agent, navitoclax, targeting the Bcl-2 family of anti-apoptotic factors. Aging Cell 15, 428–435 (2016).

6. Zhu, Y. et al. Orally-active, clinically-translatable senolytics restore α-Klotho in mice and humans. EBioMedicine 77, 103912 (2022).

7. Xu, M. et al. Senolytics improve physical function and increase lifespan in old age. Nat Med 24, 1246–1256 (2018).

8. Palmer, A. K. et al. Targeting senescent cells alleviates obesity - induced metabolic dysfunction. Aging Cell 18, e12950 (2019).

9. Farr, J. N. et al. Targeting cellular senescence prevents age-related bone loss in mice. Nat Med 23, 1072–1079 (2017).

10. Ogrodnik, M. et al. Obesity-Induced Cellular Senescence Drives Anxiety and Impairs Neurogenesis. Cell Metab 29, 1061–1077.e8 (2019).

11. Fuhrmann-Stroissnigg, H. et al. Identification of HSP90 inhibitors as a novel class of senolytics. Nat Commun 8, 422 (2017).

12. Yousefzadeh, M. J. et al. Fisetin is a senotherapeutic that extends health and lifespan. EBioMedicine 36, 18–28 (2018).

13. Zhu, Y. et al. New agents that target senescent cells: the flavone, fisetin, and the BCL- XL inhibitors, A1331852 and A1155463. Aging (Albany NY) 9, 955–963 (2017).

14. Samakkarnthai, P. et al. In vitro and in vivo effects of zoledronic acid on senescence and senescence-associated secretory phenotype markers. Aging (Albany NY) 15, 3331–3355 (2023).

15. Wissler Gerdes, E. O., Misra, A., Netto, J. M. E., Tchkonia, T. & Kirkland, J. L. Strategies for late phase preclinical and early clinical trials of senolytics. Mech Ageing Dev 200, 111591 (2021).

16. Justice, J. N. et al. A framework for selection of blood-based biomarkers for geroscience-guided clinical trials: report from the TAME Biomarkers Workgroup. Geroscience 40, 419–436 (2018).

17. Kabacik, S. et al. The relationship between epigenetic age and the hallmarks of ageing in human cells. Nat Aging (2022) doi:10.1038/s43587-022-00220-0.

18. Basisty, N. et al. A proteomic atlas of senescence-associated secretomes for aging biomarker development. PLoS Biol 18, e3000599 (2020).

19. Schafer, M. J., et al. The senescence-associated secretome as an indicator of age and medical risk. JCI Insight 5, e133668 (2020).

20. Hackl, M. et al. miR-17, miR-19b, miR-20a, and miR-106a are down-regulated in human aging. Aging Cell 9, 291–296 (2010).

21. Kocijan, R. et al. MicroRNA levels in bone and blood change during bisphosphonate and teriparatide therapy in an animal model of postmenopausal osteoporosis. Bone 131, 115104 (2020).

22. Weigl, M. et al. Longitudinal Changes of Circulating miRNAs During Bisphosphonate and Teriparatide Treatment in an Animal Model of Postmenopausal Osteoporosis. Journal of Bone and Mineral Research 36, 1131–1144 (2021).

23. Hulstaert, E. et al. Charting Extracellular Transcriptomes in The Human Biofluid RNA Atlas. Cell Rep 33, 108552 (2020).

24. Weilner, S. et al. Secreted microvesicular miR-31 inhibits osteogenic differentiation of mesenchymal stem cells. Aging Cell 15, 744–754 (2016).

25. Terlecki-Zaniewicz, L. et al. Small extracellular vesicles and their miRNA cargo are anti-apoptotic members of the senescence-associated secretory phenotype. Aging 10, 1103–1132 (2018).

26. Mensà, E. et al. Small extracellular vesicles deliver miR-21 and miR-217 as pro- senescence effectors to endothelial cells. J Extracell Vesicles 9, 1725285 (2020).

27. Jeon, O. H., et al. Senescence cell–associated extracellular vesicles serve as osteoarthritis disease and therapeutic markers. JCI Insight 4, e125019 (2019).

28. Schraml, E. & Grillari, J. From cellular senescence to age-associated diseases: the miRNA connection. Longev Healthspan 1, 10 (2012).

29. Wang, B. et al. An inducible p21-Cre mouse model to monitor and manipulate p21- highly-expressing senescent cells in vivo. Nat Aging 1, 962–973 (2021).

30. Wang, L. et al. Targeting p21Cip1 highly expressing cells in adipose tissue alleviates insulin resistance in obesity. Cell Metab 34, 75–89.e8 (2022).

31. Krishnamurthy, J. et al. Ink4a/Arf expression is a biomarker of aging. Journal of Clinical Investigation 114, 1299–1307 (2004).

32. Jeon, O. H. et al. Systemic induction of senescence in young mice after single heterochronic blood exchange. Nat Metab 4, 995–1006 (2022).

33. van Eijndhoven, M. A. J., et al. Plasma vesicle miRNAs for therapy response monitoring in Hodgkin lymphoma patients. JCI Insight 1, e89631 (2016).

34. Greussing, R. et al. Identification of microRNA-mRNA functional interactions in UVB- induced senescence of human diploid fibroblasts. BMC Genomics 14, (2013).

35. Bhaumik, D. et al. MicroRNAs miR-146a/b negatively modulate the senescence- associated inflammatory mediators IL-6 and IL-8. Aging 1, 402–411 (2009).

36. Hu, Z. et al. MicroRNA - 29 induces cellular senescence in aging muscle through multiple signaling pathways. Aging 6, 160–175 (2014).

37. Martinez, I., Cazalla, D., Almstead, L. L., Steitz, J. A. & Dimaio, D. miR-29 and miR-30 regulate B-Myb expression during cellular senescence. PNAS 108, 522–527 (2011).

38. Lessard, F. et al. Senescence-associated ribosome biogenesis defects contributes to cell cycle arrest through the Rb pathway. Nat Cell Biol 20, 789–799 (2018).

39. Zhang, X., Liu, J., Wu, L. & Hu, X. MicroRNAs of the mir-17 ∼ 92 family maintain adipose tissue macrophage homeostasis by sustaining IL-10 expression. Elife 9, 1–19 (2020).

40. Mori, M. A. et al. Role of microRNA processing in adipose tissue in stress defense and longevity. Cell Metab 16, 336–347 (2012).

41. Coppe, J.-P. et al. Senescence-Associated Secretory Phenotypes Reveal Cell- Nonautonomous Functions of Oncogenic RAS and the p53 Tumor Suppressor. PLoS Biol 6, 2853–68 (2008).

42. Navarro, F. & Lieberman, J. miR-34 and p53: New insights into a complex functional relationship. PLoS One 10, e0132767 (2015).

43. Olivieri, F. et al. MiR-146a as marker of senescence-associated pro-inflammatory status in cells involved in vascular remodelling. Age (Omaha) 35, 1157–1172 (2013).

44. Nunes, A. D. C. et al. miR-146a-5p modulates cellular senescence and apoptosis in visceral adipose tissue of long-lived Ames dwarf mice and in cultured pre-adipocytes. Geroscience 44, 503–518 (2022).

45. Rusanova, I. et al. Analysis of Plasma MicroRNAs as Predictors and Biomarkers of Aging and Frailty in Humans. Oxid Med Cell Longev (2018) 10.1155/2018/7671850.

46. Dhahbi, J. M. et al. Deep sequencing identifies circulating mouse miRNAs that are functionally implicated in manifestations of aging and responsive to calorie restriction. Aging 5, 130–141 (2013).

47. Justice, J. N. et al. Caloric Restriction Intervention Alters Specific Circulating Biomarkers of the Senescence-Associated Secretome in Middle-Aged and Older Adults With Obesity and Prediabetes in an 18-Week Randomized Controlled Trial. J Gerontol A Biol Sci Med Sci 79, 1–10 (2024).

48. Baumgart, M. et al. Age-dependent regulation of tumor-related microRNAs in the brain of the annual fish Nothobranchius furzeri. Mech Ageing Dev 133, 226–233 (2012).

49. Inukai, S., Lencastre, A. De, Turner, M. & Slack, F. Novel MicroRNAs Differentially Expressed during Aging in the Mouse Brain. PLoS One 7, e40028 (2012).

50. Somel, M. et al. MicroRNA, mRNA, and protein expression link development and aging in human and macaque brain. Genome Res 20, 1207–1218 (2010).

51. Dimri, G. P. et al. A biomarker that identifies senescent human cells in culture and in aging skin in vivo. Proc Natl Acad Sci U S A 92, 9363–9367 (1995).

52. Shannon, P., Markiel, A., Owen Ozier, Nitin S. Baliga, Jonathan T. Wang, D. R., Amin, N. & Ideker, B. S. and T. Cytoscape: A Software Environment for Integrated Models. Genome Res 13, 426 (2003).

53. Ge, S. X., Jung, D., Jung, D. & Yao, R. ShinyGO: A graphical gene-set enrichment tool for animals and plants. Bioinformatics 36, 2628–2629 (2020).

54. Diendorfer, A., Khamina, K., Pultar, M. & Hackl, M. miND (miRNA NGS Discovery pipeline): a small RNA-seq analysis pipeline and report generator for microRNA biomarker discovery studies. F1000Res 11, 233 (2022).

55. Khamina, K. et al. A MicroRNA Next-Generation-Sequencing Discovery Assay (miND) for Genome-Scale Analysis and Absolute Quantitation of Circulating MicroRNA Biomarkers. Int J Mol Sci 23, 1226 (2022).

